# Melon pan-genome and multi-parental framework for high-resolution trait dissection

**DOI:** 10.1101/2022.08.09.503186

**Authors:** Elad Oren, Asaf Dafna, Galil Tzuri, Ilan Halperin, Tal Isaacson, Meital Elkabetz, Ayala Meir, Uzi Saar, Shachar Ohali, Thuy La, Cinta Romay, Yaakov Tadmor, Arthur A Schaffer, Edward S Buckler, Roni Cohen, Joseph Burger, Amit Gur

## Abstract

Linking between genotype and phenotype is a fundamental goal in biology and requires robust data for both layers. The prominent increase in plant genome sequencing and comparisons of multiple related individuals, exposed the abundance of structural genomic variation and suggest that a single reference genome cannot represent the complete sequence diversity of a crop species, leading to the expansion of the pan-genome concept. For high-resolution forward genetics, this unprecedented access to genomic variation should be paralleled by availability and phenotypic characterization of genetic diversity, and effective integration between these layers. Here, we describe a multi-parental framework for trait dissection in melon, leveraging a novel pan-genome constructed for this crop. Melon (*Cucumis melo* L.) is an important crop from the *Cucurbitaceae* family, which display extensive phenotypic variation available for breeding. A diverse core set of 25 founder lines (*MelonCore25*) was sequenced using a combination of short and long-read technologies and their genomes were assembled *de novo*. The construction of a melon pan-genome exposed substantial variation in genome size and structure, including detection of ~300,000 structural variants and ~9 million SNPs. A half-diallel derived set of 300 F_2_ populations representing all possible *MelonCore25* parental combinations was constructed as framework for trait dissection through integration with the pan-genome. We demonstrate the potential of this unified framework for genetic analysis of various melon traits, including rind color and mottling pattern, fruit sugar content and resistance to fungal diseases. We anticipate that utilization of this integrated resource will enhance genetic dissection of important traits and accelerate melon breeding.

**Significance statement:** Pan-genomes aim to address the abundance of genome structural variation within species for improved genomic analyses. New pan-genome, constructed from *de novo* genome assemblies of 25 diverse melon (*Cucumis melo* L.) accessions is integrated with a half-diallel derived set of 300 F2 populations representing all possible parental combinations. The potential of this unified multi-parental trait dissection framework for melon genetics and breeding is presented.

## Introduction

The ability to investigate gene functions in organisms has been revolutionized in recent years by the extensive implementation of genome editing in genetic research, through CRISPR-Cas9 technology (Doudna and Charpentier, 2014). Nevertheless, genetic dissection of simple or complex traits in crop plants is still fundamentally dependent on forward genetics approaches, including the characterization of natural variation followed by genetic mapping of major Mendelian genes and quantitative trait loci (QTL). Genetic dissection of traits and QTL mapping have been accelerated by the rapid evolution of sequencing and genotyping technologies during the past two decades (Purugganan and Jackson, 2021). Parallel-sequencing-based genotyping approaches supported by multiplexing protocols (Elshire *et al*., 2011; Baird *et al*., 2008) enable high-throughput and cost-effective dense genome-wide genotyping of populations. This technology-driven advancement facilitate efficient genotypic screening of large populations and is shifting the bottleneck in mapping studies from genotyping to the phenotypic characterization of genetic diversity and the development and availability of effective diverse germplasm and segregating populations for genetic mapping within species.

Access to plant whole-genome assemblies in the early 2000s opened new avenues in crop genetics and further straightened the path from traits to QTLs and genes (Morrell *et al*., 2011). The ability to use reference genomes for physical mapping and annotation of QTL intervals has been instrumental for moving from QTLs to causative genes. An important hidden layer in the genetic variation was only recently exposed through the assembly and comparisons of multiple reference-genomes within species. The discovery of extensive structural variation (SV) led to the realization that single reference genomes does not represent the diversity within a species, and led to the expansion of the pan-genome concept (Della Coletta *et al*., 2021; Bayer *et al*., 2020).

Melons (*Cucumis melo* L., *Cucurbitaceae*) are among the most widely consumed fleshy fruits for fresh consumption worldwide (http://faostat3.fao.org/). Melons have been bred and are cultivated in nearly all of the warmer regions of the world, leading to the evolution of extensive diversity in phenotypic traits, especially in fruit traits such as size, shape, external (rind) and internal (flesh) color, sugar content, acidity, texture and aroma (Burger *et al*., 2006). This wide diversity is a source for further breeding and ongoing genetic research aimed at mapping QTLs and identifying genes affecting key horticultural and consumer-preference traits. Numerous genetic studies using diverse collections or segregating populations have focused on fruit quality traits in melon, including fruit size and shape (Martínez-Martínez *et al*., 2022; Monforte *et al*., 2014; Oren *et al*., 2020), flesh color (Cuevas *et al*., 2009; Tzuri *et al*., 2015), rind color (Feder *et al*., 2015; Oren *et al*., 2019), netting and sutures (Oren *et al*., 2020; Zhao *et al*., 2019; Diaz *et al*., 2011), sweetness and aroma (Harel-Beja *et al*., 2010; Argyris *et al*., 2017), acidity (Cohen *et al*., 2014) and ripening behavior (Oren *et al*., 2022; Pereira *et al*., 2020; Ríos *et al*., 2017). Multiple analyses have also been performed for identification and genetic characterization of resistance to pathogens (Cohen *et al*., 2022; Burger *et al*., 2003; Joobeur *et al*., 2004).

Since 2012, mapping studies in melon have benefited from the availability of a reference genome (Garcia-Mas *et al*., 2012) enabling physical mapping and annotations of QTL intervals. Resequencing of hundreds of melon accessions (S., Liu *et al*., 2020; Zhao *et al*., 2019) in recent years has further advanced the genomic analysis of candidate and causative genes. Five additional genome assemblies were published for *Cucumis melo* in the last 4 years: Three ssp. *melo* lines; the *Inodorus* line Payzawat (Zhang *et al*., 2019), the *reticulatus* line ‘Harukai’ (Yano *et al*., 2020), and the cantaloupe line ‘Charmono’ (Pichot *et al*., 2022), and two ssp. *agrestis* lines; HS (Yang *et al*., 2020) and IVF77 (Ling *et al*., 2021). These assemblies provide additional references and important information regarding the variation in melon genome structure, but a wider pan-genomic view is still lacking for this species.

In the current study, we describe an integrated multi-parental framework that we developed for genetic dissection of traits in melon. This framework is based on a representative core subset of 25 diverse melon inbred accessions that was used to construct a wide set of 300 segregating bi-parental populations derived from a 25×25 half-diallele. A novel pan-genome that we built from the sequencing and *de novo* assemblies of the 25 genomes is integrated with the multi-parental germplasm resource to create a unified trait dissection framework. We provide here examples for the effectiveness and potential of this resource for genetic analysis of various phenotypic traits in melon.

## Results and Discussion

### Selection of 25 founders from the GWAS180 diversity panel and development of the HDA25 and HDA25F_2_s populations

Our primary resource in the current project is the diverse melon collection developed at Newe-Yaar Research Center. This collection that represents the two *cucumis melo* subspecies (ssp. *melo* and ssp. *agrestis*) and 12 horticultural/varietal groups, was previously characterized for a wide range of phenotypic traits, and genotyped with ~24,000 genome-wide GBS-based SNP markers (Gur *et al*. 2017). Integration of the horticultural group classification with the phenotypic and genotypic data facilitated the informed selection of a balanced core subset composed of 25 founder lines that represent the overall diversity (hereafter *MelonCore25;* **Figure 1a and b, Table 1**; Gur *et al*. 2017). The core set includes representatives of the two cultivated sub-species and the different horticultural groups in melon as well as the broad phenotypic spectrum available for key traits, as previously described (Gur *et al*., 2017). The core set includes 17 C. *melo* ssp. *melo* lines (5 *Inodorus*, 4 *cantalupensis*, 2 *Reticulatus*, 2 *Khandalak*, 1 *Dudaim*, 1 *Flexuosus*, 1 *Ameri*, and 1 *Adzhur*), 7 lines from the *agrestis* subspecies (2 *Chinensis*, 1 *Conomon*, 3 *Makuwa*, 1 *Momordica*) and one C. *melo* ssp. *colossus* line.

**Figure 1:**
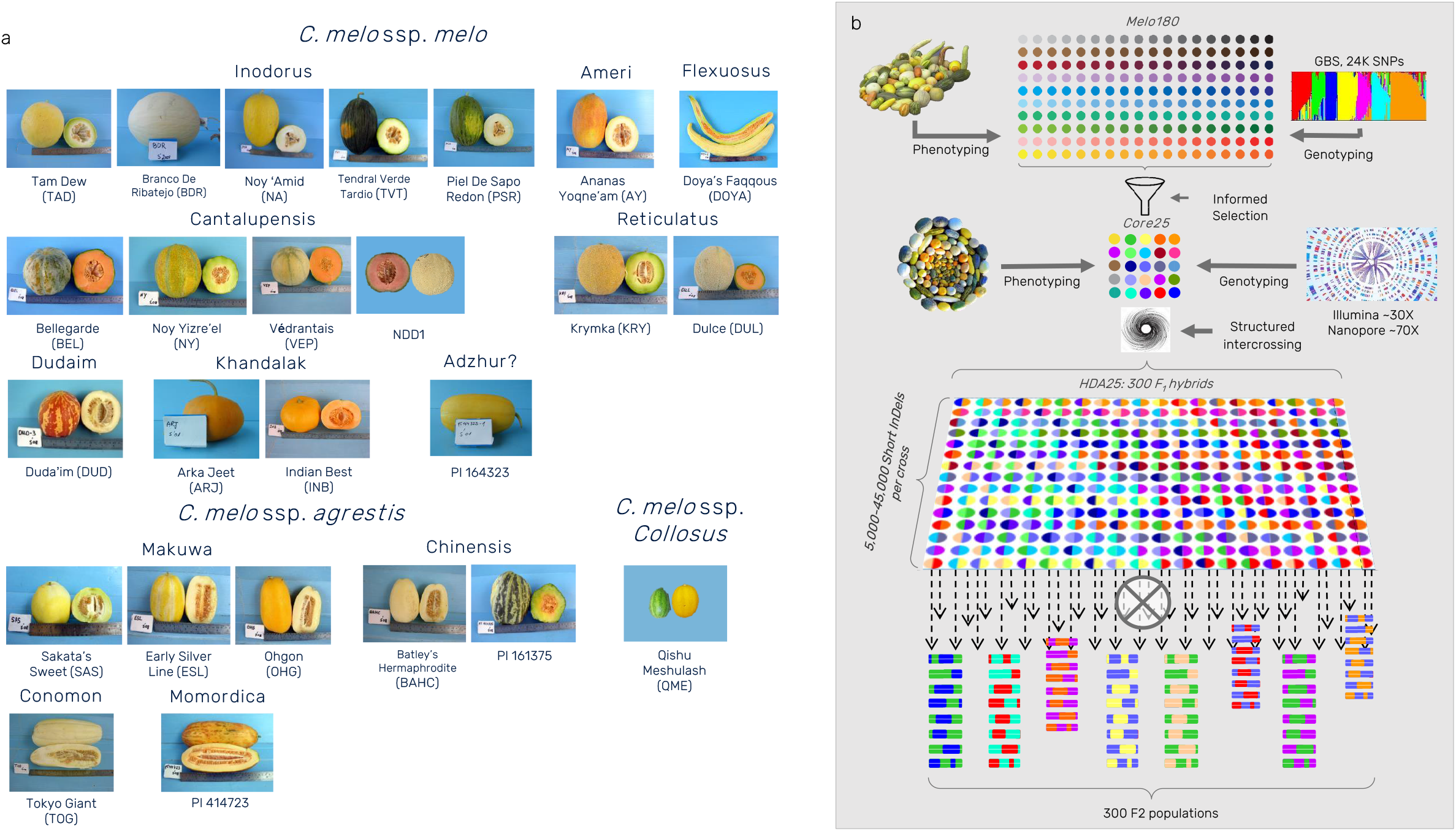
*MelonCore25* set and mapping platform development scheme. **a)** Mature fruits of the 25 founder lines, ordered by horticultural groups and sub-species division. **b)** Schematic diagram for the components and development process of the multi-parental framework.

**Table 1:**
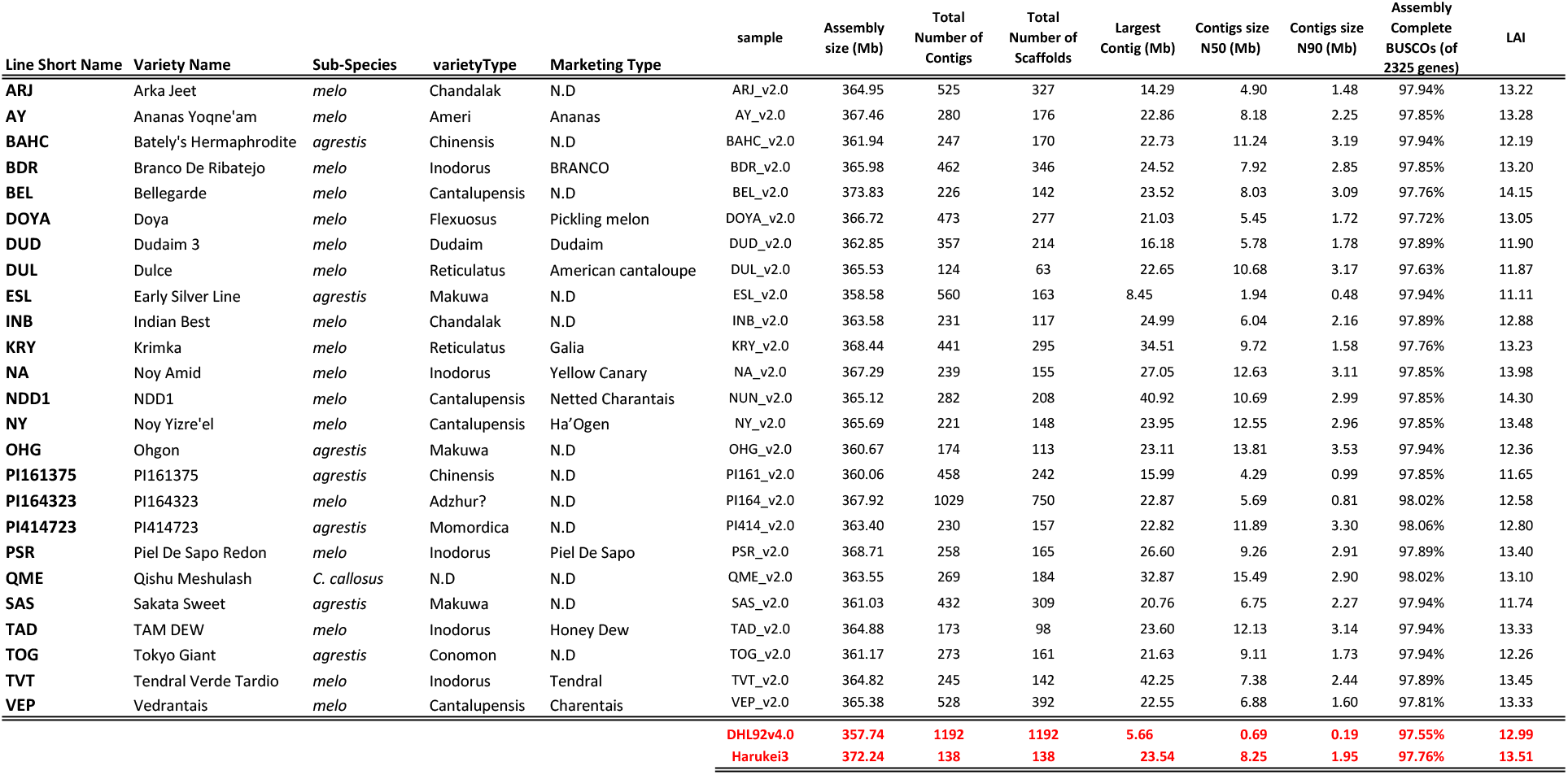
List of 25 founders (MelonCore25) and genome assembly statistics.

To fully exploit the genetic diversity of this multi-parental resource, we developed a half-dialel population by intercrossing the 25 founders in all possible combinations (*HDA25*, Dafna et al. 2021). This crossing scheme was further advanced by self-pollinations of all F1 hybrids to produce 300 distinct F_2_ populations now available for genetic dissection of melon traits (hereafter *HDA25F_2_s*, **Figure 1b**).

### Sequencing, de novo assembly of 25 chromosome-scale genomes and characterization of structural genomic variation

The 25 genomes comprising the *MelonCore25* were sequenced by Oxford Nanopore Technology (ONT) to an average depth of 70× (53-134×), generating a total sequence of 642.4 Gb with a median read length of 10Kb (4.5-24Kb) and a median read quality of 10.0 (9.2-10.6, **Sup. Table 1**). ONT reads were assembled into contigs and then polished using Illumina short reads from the previous whole-genome resequencing of these lines (average depth 30×, (Dafna *et al*., 2021)). The mean N50 of contig size across the 25 genomes is 8.4 Mb and the average number of contigs per genome is 349.5 (median 273, **Table 1**). The contigs were scaffolded into pseudomolecules through a reference-guided approach using ‘Harukei-3’ (Yano *et al*., 2020) as the reference. Assembly statistics across the *MelonCore25* expose variation in genome size ranging from 358 Mb (ESL) to 374 Mb (BEL), with a mean of 364.8 Mb (**Table 1**). These assembly statistics are comparable to the two published high-quality melon reference genomes, DHL92 v4.0 (Castanera *et al*., 2020) and ‘Harukei-3’ (Yano *et al*., 2020). Additional quality benchmarks for our assemblies are the high gene-space integrity and completeness, based on BUSCO scores that range between 97.5% to 98.1% complete genes, and the completeness of transposable elements based on the LTR Assembly Index (LAI) with an average of 12.9 (**Table 1**). Interestingly, genome size variation is correlated with the sub-species division such that genomes of ssp. *agrestis* lines are significantly smaller (avg. 361 Mb) compared to the ssp. *melo* lines (avg. 366.3 Mb**, Figure 2a, b**). Alignment of the 25 genomes and analysis of structural variation (SV) revealed extensive polymorphism in all type of SVs (InDels, inversions and translocations). Large intra-chromosomal inversions spanning across 1.6 Mb and 3.2 Mb that differentiate between most *agrestis* and *melo* lines were found in chromosomes 1 and 11, respectively (**Figure 2a**). The independent discovery of these inversions across multiple lines is another evidence for the reliability of our assemblies. We identified 108,000 large SVs (InDels and translocations) ranging from 2 Kb to 1,300 Kb (**Figure 2c**, **Sup. Table 2**). Sixty-three percent of these variants (68,500 SVs) are rare and 39,500 SVs are present in two or more lines (**Figure 2d**). We then focused on characterization of short InDels that are useful as simple PCR-based genetic markers in mapping studies. We identified 190,543 short InDel alleles in a size range of 10–2,000 bp (**Figure 2e, Sup. Table 3**). A significant proportion of these InDels represent multi-allelic polymorphisms and converge into a set of 115,802 sites. Allele frequency analysis across the 86,669 bi-allelic sites show a similar distribution pattern as in the large SVs, such that almost half of these InDels are rare (**Figure 2f**). To confirm that the *MelonCore25* comprehensively represent the genomic diversity in melon we performed pangenome analysis for the 86,669 bi-allelic InDels by plotting the number of cumulative InDels vs. number of genomes and we show that 22 lines are sufficient to capture 98% of the SV variation (**Figure 2g**). We then projected the parental short InDels profiles on the 300 *HDA25* crosses to gain a detailed description of the polymorphisms in each cross (**Sup. Table 3**). The number of polymorphic sites range between 5,082 and 45,537 sites per cross and, as expected, is strongly correlated with parental genetic distance previously calculated based on 24,000 GBS SNPs (**Figure 2h**). Short InDels coverage across the 12 melon chromosomes is relatively uniform in the *MelonCore25* pan-genome and specific crosses, regardless of the mating distance (**Figure 2i**), even in the cross between the two closely related *agrestis* lines, OHG and SAS, we identified more than 400 short InDels per chromosome. A random set of 22 short InDels from five different chromosomes were validated by PCR (**Sup. Figure 2, Sup. Table 4**).

**Figure 2:**
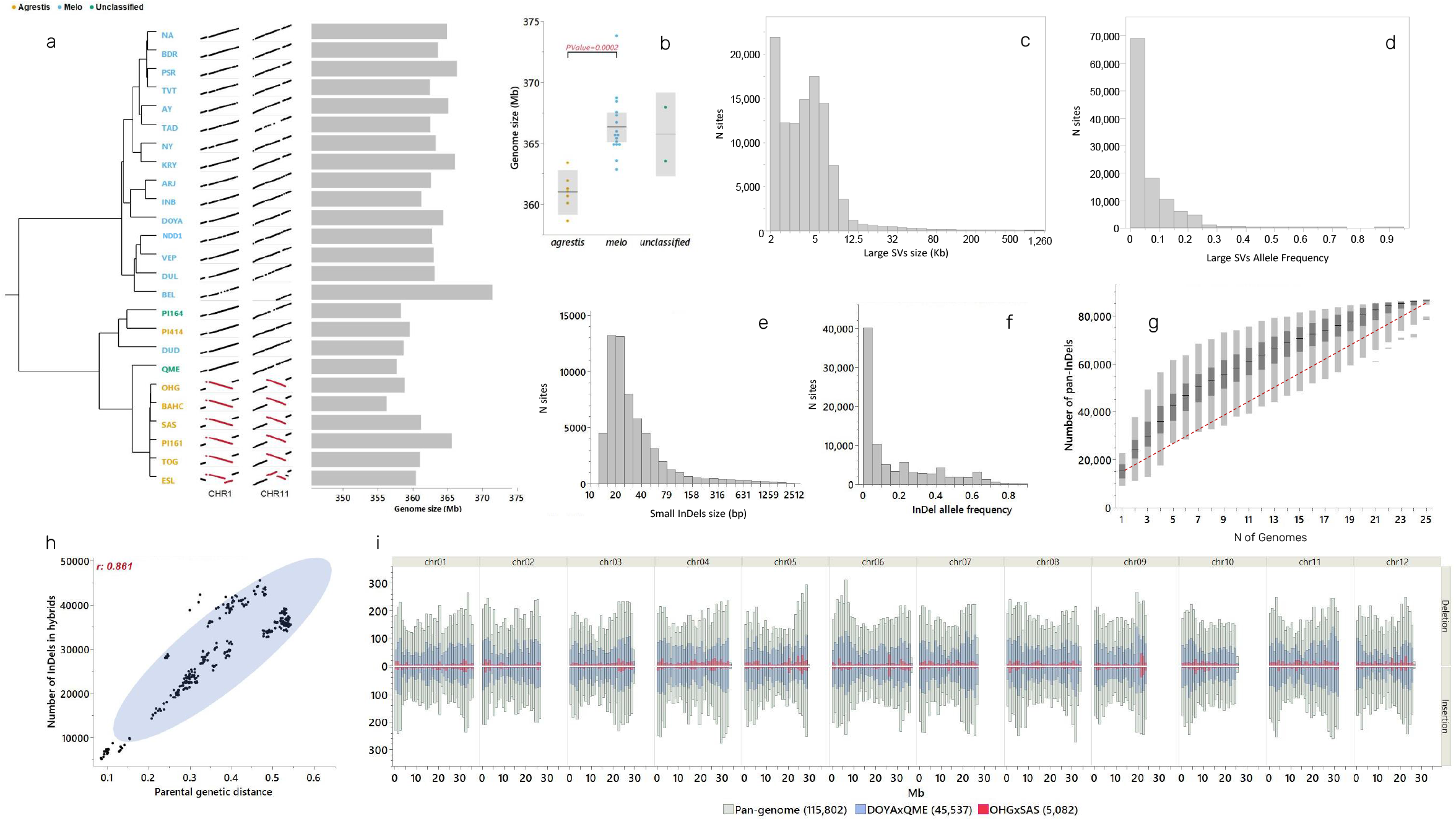
Melon pan-genome and distribution of SVs across *HDA25*. **a)** Phylogenetic tree of the 25 founder lines presented with variation in genome size and examples of large inter-chromosomal inversions on chromosomes 1 and 11 in ssp. *agrestis* lines. **b)** Genomes of ssp. *agrestis* lines are significantly smaller than ssp. *melo* lines. **c)** Size distribution of 108,011 large SVs (>2 Kb). X axis is presented in log scale. **d**) Distribution of allele frequency of 108,011 large SVs (>2 Kb). **e**) Size distribution of 86,669 short (<2 Kb) bi-allelic InDels across MelonCore25. X axis is presented in log scale. **f)** Distribution of allele frequency across 86,669 bi-allelic sites (<2 Kb). **g**) Pan InDels analysis. Horizontal black lines are the mean number of pan InDels under corresponding number of genomes. Dark boxes are the 50 percentile and light boxes are 99 percentile **h)** Correlation between parental distance (calculated using 24,000 genome-wide GBS SNPs) and number of polymorphic short InDels across 300 *HDA25* crosses. **i)** Number of polymorphic short InDels by chromosomes and bins, in the pan-genome and two crosses. In parentheses is the total number of short Indels for each group.

Genome sequences are available to date for 18 different cucurbit species revealing the variation in genome size and organization across the *Cucerbitaceae* family and reflect the evolution of this highly diverse group (Ma *et al*., 2022). Information on intra-specific variation in genome size within this family is, however, still relatively limited. Genomes of 11 *cucumis sativus* accessions were recently sequenced and compared, exposing difference of 26 Mb (11%) between the largest and smallest genomes, as well as 56,214 SVs (Li *et al*., 2022). We found here size variation of 16 Mb (4.5%) between the largest and smallest genomes across *MelonCore25*. The significant difference between the *agrestis* and *melo* sub species support the validity of these differences and imply on the possible evolutionary significance of within-species genome size variation (Biémont, 2008).

### Workflow for utilization of the multi-parental framework

Our integrated framework relies on the connection of the structured multi-parental set of populations (*HDA25F_2_s*) with phenotypic data, whole-genome assemblies and derived variome of the 25 parental lines (**Sup. Tables 2 and 3**). The proposed utilization workflow is described in **Sup. Figure 1**: briefly, a trait of interest should first be scanned across the *MelonCore25* using the database, for existing trait, or through direct phenotypic analysis, for new trait. Following reliable phenotyping, we found that projecting the phenotypic data on the genetic PCA of *MelonCore25* is a useful way to integrate genotype and phenotype for informed selection of parental lines differing in the trait of interest and reflect varying parental genetic distance. Following the selection of parental lines, optimal F_2_ population can be obtained from *HDA25F_2_s* set for characterization of mode of inheritance and genetic mapping. Depending on the genetic architecture and complexity of the trait, the relevant mapping strategy should be applied, and accordingly the population size and generation should be adjusted. Parental genome assemblies of the selected population are available as reference for Bulk-sequencing analyses (BSA-Seq, QTL-Seq) and for markers development for further fine-mapping. Detailed annotation of simple or quantitative trait locus (QTL) intervals can also benefit from referring to the specific parental genomes. After the initial whole-genome mapping, the *HDA25F_2_s* set is a valuable resource for selection of additional relevant crosses for QTL validation or allelism testing. Genomes of the 25 founders can then be used for mining allelic variation at candidate genes.

### Examples for implementation

#### Genetics of pigment intensity and mottled rind

Mottled rind is a common characteristic in melon and is distributed and expressed in different patterns across our diverse *GWAS180* collection (**Figure 3a**). As with other traits, *MelonCore25* capture this diversity with parallel proportions of mottled and smooth rind, as in the wider *GWAS180* collection (**Figure 3b**). To present a comprehensive description of the genetic architecture of this trait, we projected the rind-mottling trait on the genetic PCA plot and selected several mottled and smooth lines for focused genetic analyses (**Figure 3c**). We started by mapping mottled rind in the cross between ‘Dulce’ (DUL, *C. melo var. reticulatus*) that has uniform dark rind, and the mottled line PI414723 (PI414, Indian phut snapmelon, *C. agrestis, Momordica* group), using a previously described RILs population (**Figure 3d**, (Galpaz *et al*., 2018)). Whole-genome linkage analysis resulted in the mapping of a single major trait locus to ~400Kb interval on chromosome 2 (**Figure 3e**). We then zoomed in on this locus and validated its association with mottled rind through independent GWA analysis of mottled rind trait on our melon diverse collection (**Figure 3f**, n=177, (Gur *et al*., 2017)). The integration of linkage and association analyses allowed us to narrow the mapping to a common genomic interval of less than 200Kb, which we further confirmed through substitution mapping using detailed analysis of recombinants on the PI414×DUL RILs (**Figure 3g**).

**Figure 3:**
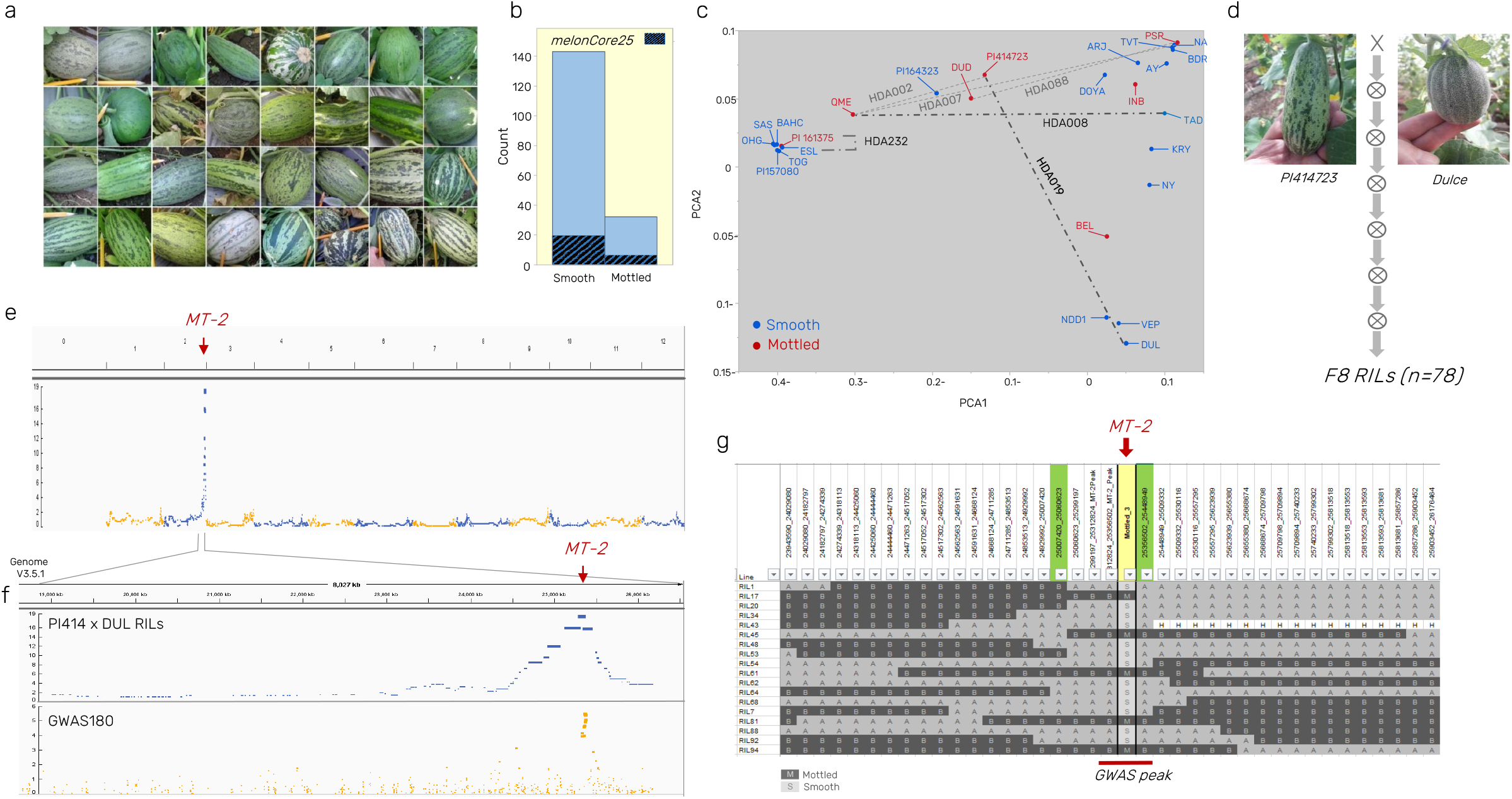
Genetic analysis and mapping of mottled rind. **a)** Representative mottled rind fruits from the *GWAS180* collection. **b**) Parallel frequencies of smooth and mottled rind lines in the *GWAS180collection* and the *MelonCore25* set. **c)** Projection of rind pattern (smooth or mottled) on the genetic PCA plot of the *MelonCore25* set. Dashed lines represent crosses (and derived segregating populations) used for genetic analysis of mottled rind trait. **d)** PI414 and DUL, parental lines of the RILs population used for mapping the mottled rind trait. **e)** Linkage mapping results of the *MT-2* locus to chromosome 2. **f)** Zoom in on chromosome 2 mapping in the PI414×DUL RILs and comparison to Manhattan plot from GWA analysis of the *GWAS180* collection. **g)** Substitution mapping validation for the *MT-2* locus using recombinants from the PI414723×DUL RILs. Green cells represent flanking markers for the interval. Yellow cell is the rind phenotype column. A= DUL allele, B=PI414 allele. GWAS peak interval is represented by the red horizontal line.

To expand the analysis of mode of inheritance of the mottled rind trait, we analyzed rind pattern of young fruits across the *HDA20* population (subset of 20 founder lines from *MelonCore25*, and their 190 half-diallel F1 hybrids). Results were consistent with a single dominant gene inheritance across this wide array, as all crosses with mottled parent resulted in mottled F1, and smooth hybrids were obtained only by the combination of two smooth parents (**Figure 4a).** We then selected two additional segregating populations from crosses between mottled and smooth lines: 1) F_2_ population from the cross between the mottled *Cucumis colossus* accession QME and the smooth honeydew line, TAD (**Figure 4b**), and 2) F_2_ population from the cross between two closely related *agrestis* lines, the mottled line PI161375 (group *Chinensis*) and the light and smooth rind line ESL (Early Silver Line, group *Makuwa*) (**Figure 4c**).

**Figure 4:**
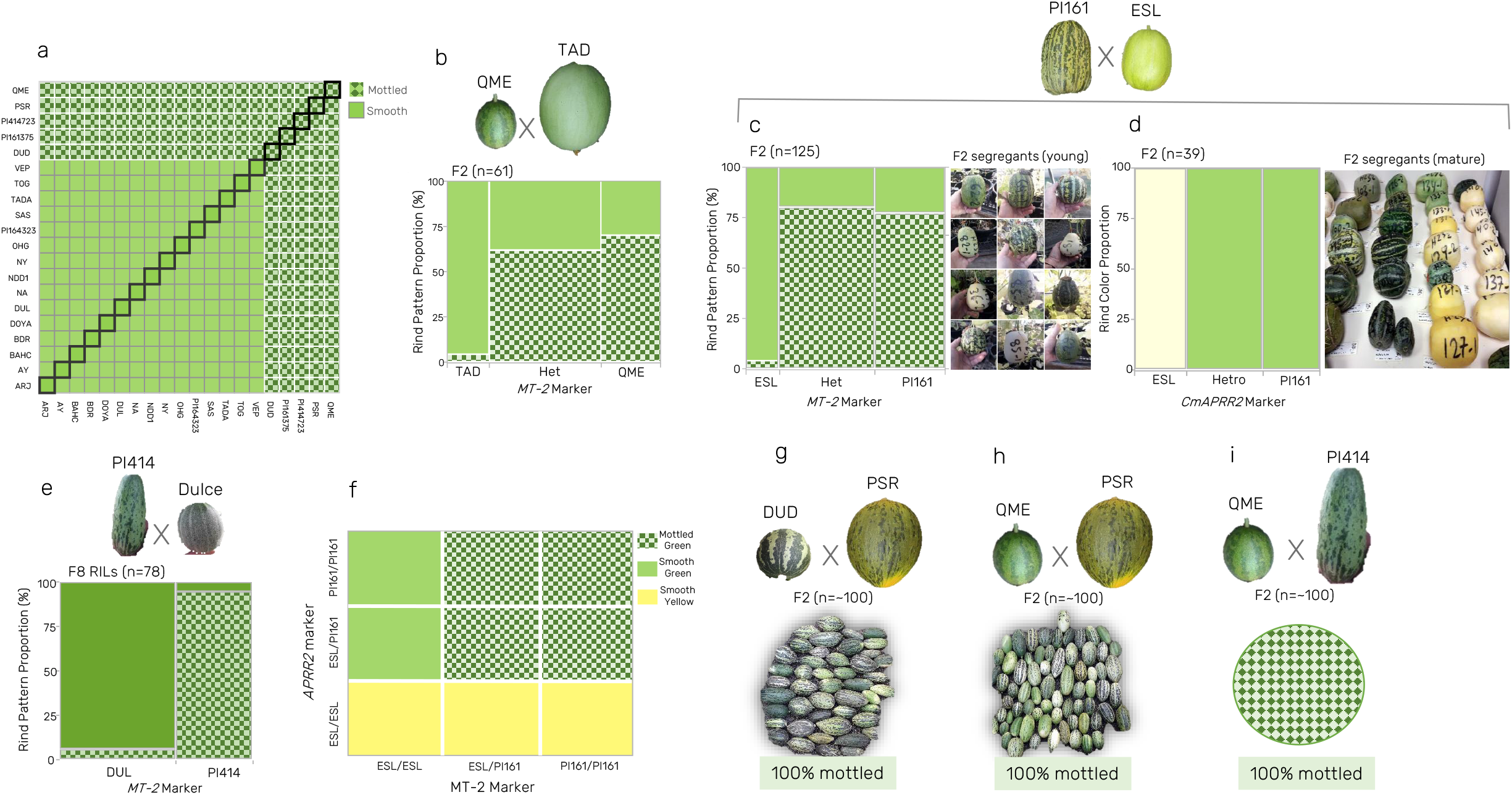
Multi-parental characterization of mottled rind trait. **a)** Inheritance of rind pattern across *HDA20* set (190 half-diallel F1s and their 20 parents). Each cell represent the combination of parents from the horizontal and vertical axes. Diagonal represent the inbred parents’ phenotype. **b)** Contingency analysis for the segregation of rind pattern against *MT-2* marker in the QME×TAD F2. Het=Heterozygote. **c)** Contingency analysis for the segregation of rind pattern against *MT-2* marker in the PI161375×ESL F2. Het=Heterozygote. Examples for young fruit segregants (smooth and mottled) are displayed. **d)** Contingency analysis for the segregation of rind color against *APRR2* marker in the PI161375×ESL F_2_. Het=Heterozygote. Examples for mature fruit segregants (light and dark rind) are displayed. **e)** Contingency analysis for the segregation of rind pattern against *MT-2* marker in the PI414723×DUL RILs. **f)** Schematic pivot table that present the co-segregation of *APRR2* and *MT-2* markers against rind color and pattern in the PI161375×ESL F_2_. **g)** Allelism test for the mottled rind trait in the DUD×PSR F2. **h)** Allelism test for the mottled rind trait in the QME×PSR F_2_. **i)** Allelism test for the mottled rind trait in the QME×PI414723 F_2_.

In both populations rind mottling was phenotyped on young fruits (10-15 days post anthesis) and we found significant association between markers that we developed at the *MT-2* locus and mottled rind segregation (**Figure 4b, c**). Interestingly, segregation and association patterns in these populations were different compared with the PI414×DUL RILs population (**Figure 4e**), such that in both crosses, while groups homozygous to the smooth parent allele at the *MT-2* marker were perfectly associated with smooth rind, the heterozygotes and homozygous for the mottled parent genotypes included also significant proportion of smooth rind segregants (**Figure 4 b, c, e**). Such pattern is indicative of possible epistasis with another locus affecting this trait. Recently, Shen *et al* proposed *CmAPRR2* as epistatic over *MT-2* in controlling mottled rind phenotype in a cross between a mottled *agrestis* inbred (Songwhan Charmi, SC) and a smooth *ssp. melo* line (Mi Gua, MG) (Shen *et al*., 2021). The PI161375×ESL population displayed very clear segregation to light and dark rind color and was therefore also used to test the association between *CmAPRR2* gene and rind pigment intensity in mature fruits. In a previous study, we proposed that the light rind in ESL is a result of single base insertion in exon9 of the *CmAPRR2* gene causing a frameshift that lead to major modification in predicted protein sequence (Oren *et al*., 2019). PI161375 carry the normal (‘dark’) allele in this gene and indeed we found complete co-segregation of the SNP in *CmAPRR2* and rind color, supporting the proposed exon9 polymorphism as causative (**Figure 4d**). We combined the genotypic data of *CmAPRR2* marker and the *MT-2* marker in the ESL×PI161375 population and confirmed that the segregation of the two genes perfectly explain the smooth and mottled rind segregation (**Figure 4f**), as the light allele of *CmAPRR2* is epistatic over *MT-2*.

To further validate the assumed allelism between different mottled rind sources, we tested three additional crosses between different mottled lines: 1) Cross between DUD, a line from *Dudaim* group and PSR, a Piel de Sapo line from the *inodorous* group (**Figure 4g**). 2) Cross between QME and PSR (**Figure 4h**), and 3) Cross between QME and PI414 (**Figure 4i**). In all three crosses, 100% of the F2 segregants (n=~100) displayed mottled rind fruits, providing another supportive evidence that the different mottled rind patterns across melon diversity are allelic and caused by the *MT-2* gene.

Rind color and pigmentation pattern are important attributes of melon fruit appearance. We previously reported *CmAPRR2* gene as a key regulator of rind and flesh pigment intensity in melon (Oren *et al*., 2019). That study benefitted from integrative analysis of pigment variation across multi-parental germplasm. Genomes resequencing of *MelonCore25* were used for detailed analysis of haplotypic variation in the *cmAPRR2* gene and allowed detection of multiple independent predicted causative mutations across the diversity that were supported by comprehensive allelism tests using the *HDA25* population. This high-resolution multi-parental analysis exposed the complex haplotypic variation in the *CmAPRR2* gene and resolved the unexpected lack of signal in GWAS for this conserved central pigment accumulation regulator (Oren *et al*., 2019). Co-segregation analysis between light rind phenotype and ESL allele in the current study provides another support for the multi-allelic model and predicted causality of an exon9 point mutation in *cmAPRR2* for the light rind of this line (**Figure 4d**).

We describe here the mapping of mottled rind trait using a combination of segregating population and diverse collection (**Figure 3 e, f**). We mapped the trait to a ~115 Kb interval with 14 annotated genes. Our mapping results coincide with previous mapping of the *Mt-2* gene by parallel analysis of multiple crosses to support the allelic nature of mottled rind trait from different sources. Mapping of mottled rind trait to the *MT-2* locus on chromosome 2 was reported in three previous studies conducted using different parental lines. Lv *et al*., (2018) mapped the spotted rind trait in var. *chinensis* background to 280 Kb interval on chromosome 2. Pereira *et al*., (2018) described the mapping of mottled rind to the same genomic region in a cross between the smooth rind “Védrantais” (ssp. *melo, cantalupensis* group) and mottled “Piel de Sapo” line (ssp. *melo, inodorus* group). Recently, Shen *et al*., (2021) mapped mottled rind to a 40.6 Kb within the same interval harboring 6 genes, using a ssp*. agrestis* var*. chinensis* line as the trait donor. Conclusive evidence for the causative gene and sequence variants is yet to be provided. The mapping interval reported in our study overlap with the 40.6 Kb and the detailed genomic variation available now for *MelonCore25* can assist in identifying such causative variation. We analyzed SNPs and SVs within the 6 candidate genes in the 40.6 Kb interval (**Sup. Tables 5 and 6**). This region possess 62 exonic SNPs, 21 of them are non-synonymous polymorphisms, 14 are UTR polymorphisms and 27 are synonymous or splice-site polymorphisms. Interestingly, none of these polymorphisms display significant association with the mottled rind phenotype across the *MelonCore25* set. The broad multi-parental analyses here and in previous studies support the notion that mottled rind variation is determined by a single gene across melon diversity which could suggest a possible monophyletic nature of this trait. However, the lack of association of exonic polymorphism across the 25 founder lines can suggest that the variation is either regulated on other levels (expression, translation, etc.) or by multi-allelic variation caused by independent mutations in the causative gene, as we previously descried for the APRR2 gene in melon (Oren *et al*., 2019). Resolution of this will be achieved once the causative sequence variation will be discovered and characterized.

#### Genetics of sugar accumulation

Fruit sweetness is the primary parameter defining melon quality and consumer preference. Multiple studies described the physiology and complex genetic architecture of sugars accumulation and fruit sweetness (Dai *et al*., 2011; Harel-Beja *et al*., 2010; Schaffer *et al*., 1987; Leida *et al*., 2015; Argyris *et al*., 2017). The *MelonCore25* was characterized for total soluble solids (TSS, °Bx) content of mature fruits grown in replicated open-field experiments. Mean TSS of the lines across this core set varied from 4 to 15 °Bx (**Figure 5a**). This variation is parallel to the range displayed across our broader *GWAS180* collection (Gur *et al*., 2017) and reflects the wide spectrum of fruit sweetness within *Cucumis melo* and the relevance of studying the genetics and physiology of sugar transport and accumulation within this species. Projection of TSS values on genetic PCA of *MelonCore25* set (**Figure 5b**) show that fruit sweetness is not independent of population structure, as most of the ssp. *melo* lines are sweet, with the exception of non-sweet lines from the *Flexuosus* group, while among the cluster of the ssp. *agrestis* lines in our core set, there is representation of high and low TSS. Based on this integrated view, we selected several crosses between high and low TSS lines that represent different combinations of groups and sub-species classifications. The rationale is to characterize the genetic architecture of sugar accumulation in multiple crosses that represent melon diversity, which will facilitate comprehensive view of the genetic architecture of fruit sweetness. For example, *HDA232* is a cross between two closely related *agrestis* lines differing in TSS: Early Silver Line (ESL, *Makuwa* group, Bx=14.6) and PI161375 (*Chinensis* group, Bx=7.0), and therefore such a cross potentially represents allelic variation in a small number of major sugar QTLs; *HDA243* is an intra sub-specific combination between genetically distant lines, the sweet *agrestis* line, Sakata Sweet (SAS, *Makuwa* group, Bx=14.0) and the non-sweet *melo* line DOYA (*Flexuosus* group, Bx=3.8), and is therefore expected to reflect variation in multiple sugar QTLs across the genome. Examples of three selected segregating populations that were advanced for mapping sugars QTLs are provided: in **Figures 5d-f.** We show the TSS correlation between F3 and F4 family means in the cross between the sweet honeydew line, TAD (Bx=15.2) and the non-sweet *Cucumis colossus* accession QME (Bx=8.0). Tail segregants from 208 F3:F4 families are highlighted for QTL-Seq analysis (**Figure 5d**). **Figure 5e** shows the TSS correlation between F4 and F5 family means in the cross between TAD and a different non-sweet line, PI164323 (Bx=4.9). Tails segregants from this population were also selected for mapping the main sugar QTLs in this cross. **Figure 5f** presents the variation in TSS and tails segregants selection in the F5 population derived from the inter-sub-specific cross between the sweet *agrestis* line, Sakata Sweet (SAS) and the non-sweet ssp. *melo flexuosus* line DOYA.

**Figure 5:**
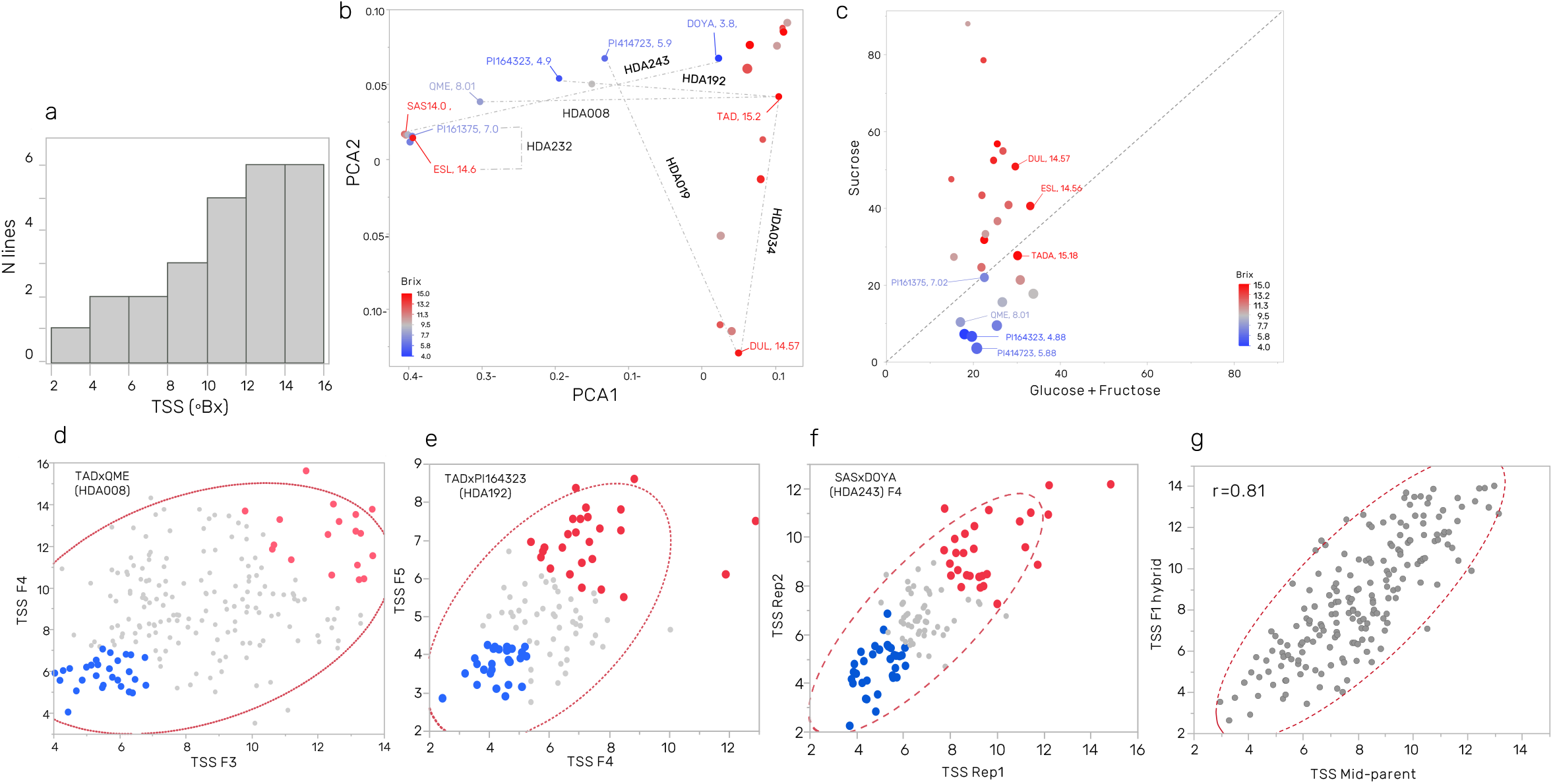
Variation in fruit sugars across *MelonCore25* and selected segregating populations for genetic mapping. **a)** Distribution of TSS of mature fruits across *MelonCore25*. **b)** Projection of lines TSS on the genetic PCA. Dashed lines represent crosses (and derived segregating populations) used for genetic dissection of fruit sugars accumulation. **c)** Correlation between monosaccharides (Glucose and Fructose) and disaccharide (Sucrose) content in mature fruits of *MelonCore25*. **d)** Selection of tails segregants through the analysis of correlation between F3 and F4 lines in the TAD×QME (*HDA008*) cross. High and low TSS tails are represented in red and blue, respectively. **e)** Selection of tails segregants through the analysis of correlation between F4 and F5 lines in the TAD×PI164323 (*HDA192*) cross. High and low TSS tails are represented in red and blue, respectively. **f)** Selection of tails segregants through the analysis of correlation between replications in the SAS×DOYA (*HDA243*) F4 population. High and low TSS tails are represented in red and blue, respectively. **g)** Correlation between TSS of parental mean (mid-parent) and TSS of their F1 hybrid across 190 F1s (*HDA20* population).

To expand our view on the inheritance of sugar accumulation using the multi-parental framework, we analyzed TSS in the *HDA20* sub-population composed of 20 parents from *MelonCore25* alongside their 190 half-diallel F1 hybrids. In order to gain broad insight on the mode-of-inheritance of fruit TSS, each of the 190 F1s was compared to its parental lines. We find all modes of inheritance across the broad *HDA20* set (**Sup. Figure 3**); however, the highly significant correlation (*r*=0.81) between mid-parent and F1s TSS (**Figure 5g**) across the 190 diverse hybrid groups is indicative of the prominent additive mode of inheritance of TSS through melon diversity. Taken together with the complex genetic architecture of fruit TSS, these results imply that the use of multiple bi-parental populations could be an effective path to genetically dissect this trait and develop marker–assisted selection protocols to breed sweet melons.

In addition to TSS measurements, *MelonCore25* was also analyzed for sugars composition. As previously shown (Burger and Schaffer, 2007; Schaffer *et al*., 1987), the sugar content continuum in melon is explained by the variation in sucrose rather than in its hexoses building blocks—glucose and fructose (**Figure 5c**). This unidirectional variation, reflected through the bivariate view of the hexoses versus sucrose content across *MelonCore25* (**Figure 5c**) is expressing the variation in sugars composition, and the variable proportion of sucrose from the total sugars in melon—the sweet lines contain 85% sucrose and the non-sweet only 15%. This variation is a very effective substrate for establishing genetic studies focused on sugars metabolism in melon fruit.

While there is a very clear distinction between sweet and non-sweet melon types, previous studies suggest that the genetics of sucrose metabolism and accumulation is complex and most likely controlled by multiple QTLs with small effects rather than variation in major genes (Diaz *et al*., 2011; Leida *et al*., 2015). The physiological and biochemical characterization of sugars metabolism performed thus far in melon provide many candidate genes related to sugars metabolic pathway and transport (Dai *et al*., 2011), but causative genes explaining the phenotypic variation are yet to be discovered through effective genetic analyses. Argyris *et al*. (2017) analyzed multiple segregating populations from two different crosses between sweet and non-sweet parents for mapping sugars accumulation QTLs. We expand this approach by targeting multiple crosses from the *HDA25F_2_s* set. Our core set represent the whole sweetness spectrum in melon, from Brix 4 to Brix 15, and we selected several crosses between high and low-sugar parents representing different genetic combinations (**Figure 5b**). Using the genome assemblies of all the participating parental lines in these crosses, we will perform parallel QTL-Seq analyses using tail segregants (**Figure 5 d-f**) and anticipate efficient annotations of sugar accumulation QTL intervals in these populations. In addition to the complete representation of SNP and short InDels variation in these populations, a detailed comparative analysis of structural variation within QTL intervals or presence-absence variation (PAV) analysis of sugar metabolism and transport genes can be further analyzed and associated with phenotypic variation.

#### Genetics of resistance to soil-borne pathogens

##### Variation in resistance to Fusarium wilt and haplotypic variation in Fom-2 gene across MelonCore25

Fusarium wilt is an important disease affecting melon production, caused by the soil-borne fungus *Fusarium oxysporum* f sp. *melonis (F. o. melonis*). We characterized the response of young seedlings of the *MelonCore25* set to *F. o. melonis* races 1 and 2 inoculation. For both pathogen races the whole spectrum of responses from resistant to susceptible is represented across this core set. Projection of the disease symptoms indexes on the genetic PCA display the division of the response to race 1 by subspecies, such that the ssp. *agrestis* lines are resistant and ssp. *melo* lines are all intermediate or highly susceptible (except for VEP). It also confirms the expected differential response of the lines to races 1 and 2 (**Figure 6a** and **b**). This differential response is most prominent among the six *agrestis* lines that are resistant to race 1 and highly susceptible to race 2 (**Figure 6a** and **b**). A major resistance gene (*Fom-2*) conferring resistance to *F. o. melonis* races 0 and 1 was previously mapped and positionally cloned (Joobeur *et al*., 2004). Genetic marker was further developed within the *Fom-2* gene for marker assisted selection in melon breeding (Wang *et al*., 2011). We used our multi-parental platform to evaluate cases of limited predictive value of *Fom-2* marker across different breeding germplasm. The 25 genome assemblies allowed us to obtain detailed alignment of the *Fom-2* (Melo3C021831) gene sequence and detect multiple polymorphisms, including large InDel between exon2 and exon3 (**Figure 6c**). In total, we found 45 polymorphic sites within exons in the *Fom-2* gene, 26 of them are non-synonymous polymorphisms resulting in different types of effect on expected protein sequence (**Sup. Table 7**). Interestingly, this extensive sequence diversity is translated into only 3 main haplotypes across the *MelonCore25* set: Haplotype 1 is specific and common to the ssp. *agrestis* lines in our core set and is similar to the DHL92 reference in this region (**Figure 6c, d**). Haplotype 2 is distributed across the different groups within the ssp. *melo* lines and characterized by multiple SNPs along the gene that are part of linkage disequilibrium (LD) block. Haplotype 3 is common to ssp. *melo* lines from different groups and contain an insertion of ~1,100 bp compared to the reference, and few additional SNPs in LD (**Figure 6c, d**). While we found significant association between these *Fom-2* haplotypes and Fusarium race 1 resistance across *MelonCore25* set, none of the polymorphisms independently explain the variation (**Figure 6**). The only exception for this haplotypic association is the ssp. *melo* line VEP (*Charentais* group, **Figure 6c**: Haplotype 2, resistant to race 1), suggesting additional genes involved. The cross between VEP and the related susceptible line DUL (*Reticulatus* group, Haplotype 2) is therefore an informed selection of potential population for mapping these other genetic loci (**Figure 6a**).

**Figure 6:**
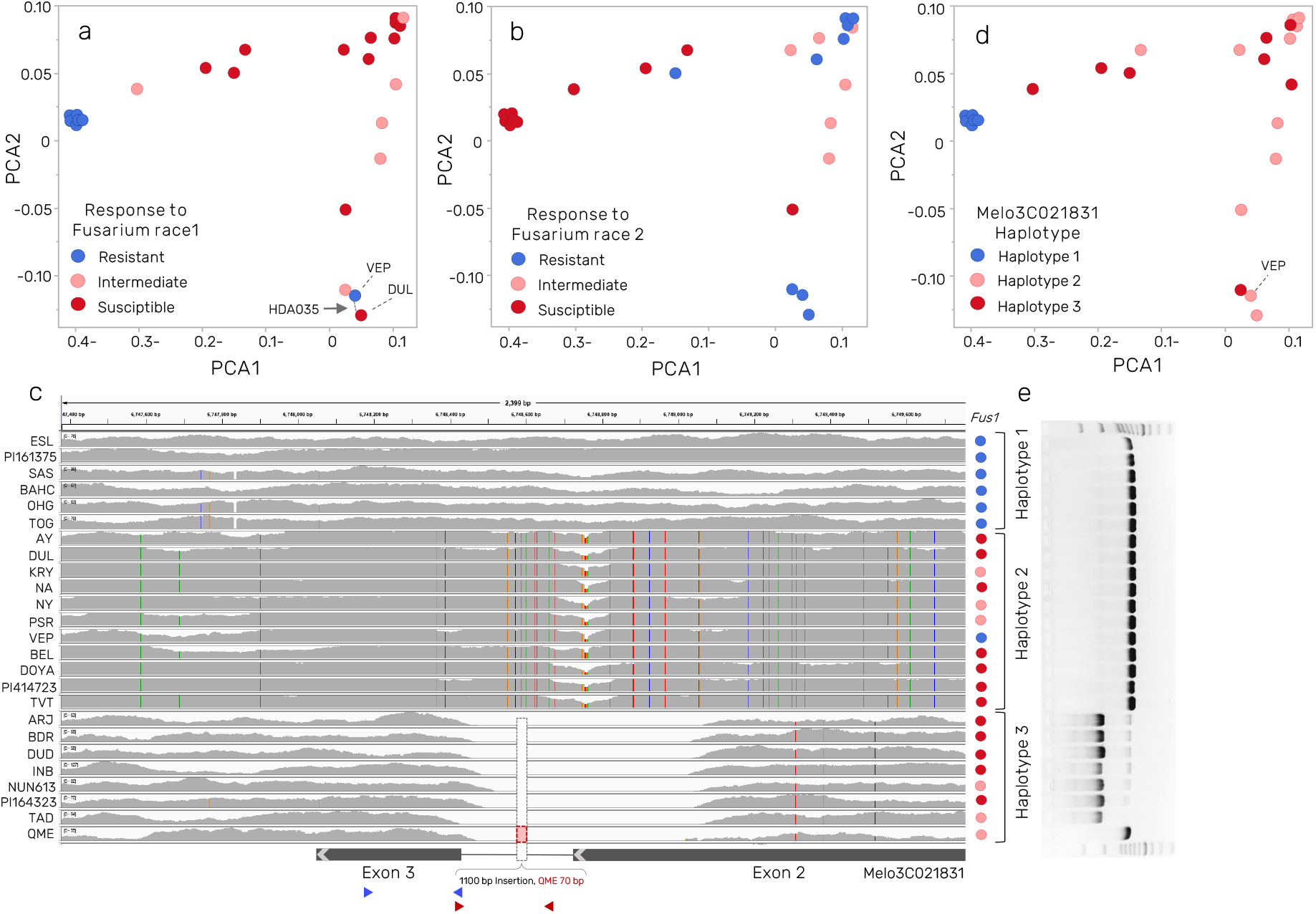
Variation in response to *Fusarium oxysporum* f.sp. *melonis* races 1 and 2 and haplotypic variation in *FOM-2* gene across *MelonCore25.* **a)** Projection of lines response to Fusarium race 1 on the genetic PCA. **b)** Projection of lines response to Fusarium race 2 on the genetic PCA. **c)** Haplotypic variation in the *FOM-2* gene (MELO3C021831) across *MelonCore25*. Vertical colored lines are SNPs. Gray histograms reflect short read depth. Dashed rectangle in haplotype 3 is the1,100 bp insertion discovered through *Nanopore* sequencing and *de novo* assemblies. Remarkably, a ~500 bp deletion was initially reflected in this region based on absence of Ilumina short read alignments. The actual insertion was discovered only based on the long-reads assemblies (**Sup. Figure 5**), and was confirmed by PCR, presented in **Figure 6e**). Colored dots correspond to the Fusarium race 1 response as presented in **Figure 6a**. Part of the gene model is presented below the haplotypic view. Triangles bellow the gene model are PCR primers for FOM-2 CAPs markers. Blue, marker from (Oumouloud and Otmani, 2015). Red, CAPs marker developed for the Newe-Yaar breeding program (**Sup. Table 8**). **d)** Projection of lines *FOM-2* haplotype on the genetic PCA. **e**) Gel image of PCR validation of the 1,100 bp InDel.

##### *Genetic analysis of resistance to Fusarium oxysporum* f. sp. *radicis-cucumerinum (FORC)*

Fusarium root and stem rot caused by the fungus *Fusarium oxysporum f. sp. radicis-cucumerinum* (FORC) is a disease in greenhouse cucumbers and melons (Punja and Parker, 2000; Vakalounakis *et al*., 2005). Genetic variation in the response to FORC inoculation was previously evaluated (Cohen *et al*., 2015; Elkabetz *et al*., 2016). We characterized the response to FORC across *MelonCore25* set and the whole spectrum from susceptibility to resistance is displayed (**Figure 7a**). Projection of disease severity index (DSI) on the genetic PCA plot exposes that the level of resistance to this pathogen is independent of population structure (**Figure 7b**). Multiple potential crosses between resistant and susceptible lines are available at the *HDA25F_2_s* set, and we used the RILs population developed from the cross between TAD (*Inodorous* group, intermediate resistance) and DUL (*Reticulatus* group, susceptible) (Oren *et al*., 2019) to genetically dissect this trait. The RILs population displayed transgressive segregation in the response to FORC inoculation with lines more resistant than TAD or more susceptible than DUL (**Figure 7c**). This population was genotyped with 89,343 GBS SNPs defining 2853 recombination bins (Oren *et al*., 2019) and through whole-genome QTL mapping analysis we identified a major QTL on chromosome 7 (**Figure 7d**) that explain 22% of the FORC DSI variation (*p*=2×10^-8^, **Figure 7e**). We used the parental *de novo* assembled genomes and derived melon variome database to extract informative InDel markers for further backcrossing for validation and fine-mapping of this QTL.

**Figure 7:**
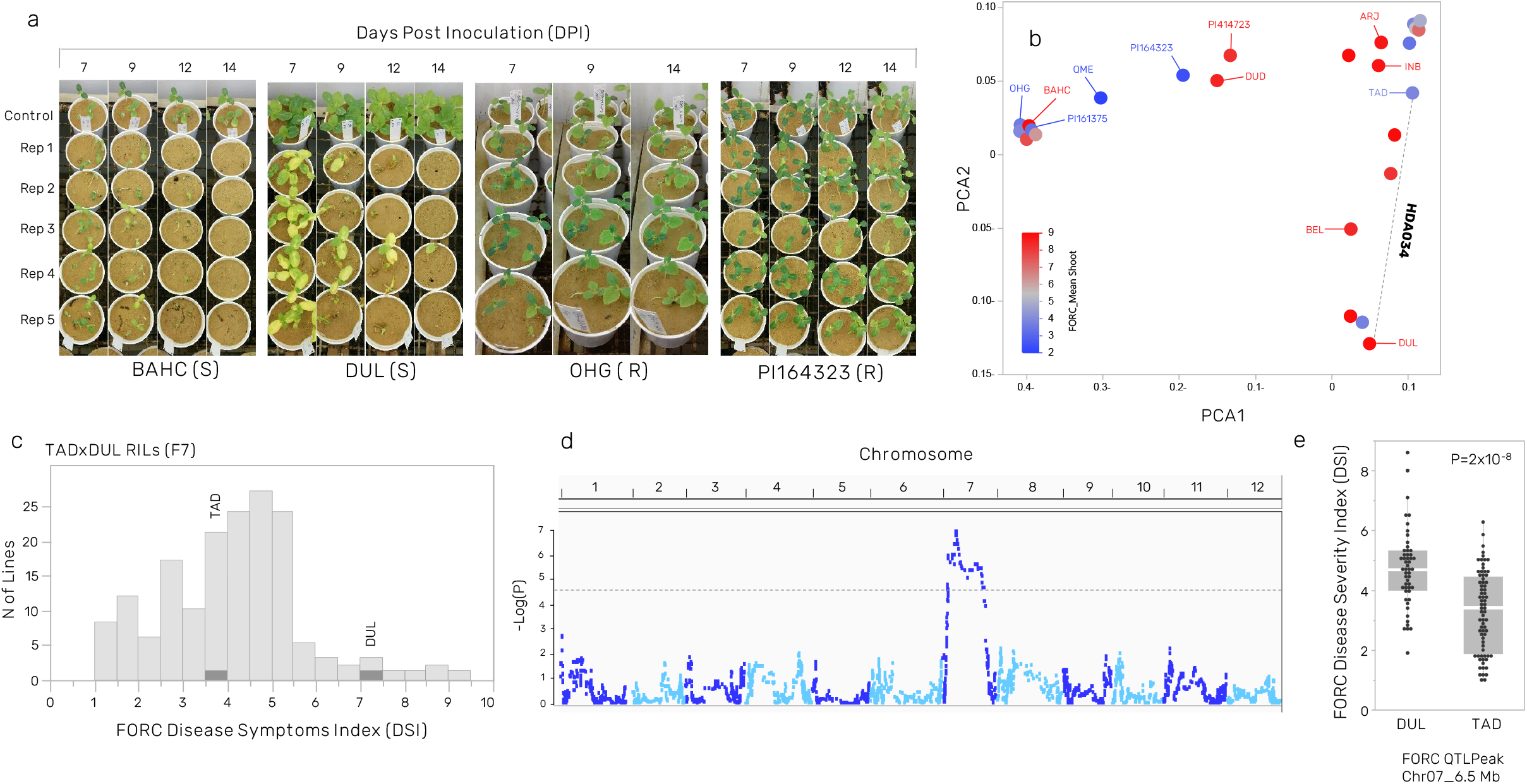
Variation in response to *Fusarium oxysporum* f. sp. *radicis-cucumerinum* (FORC) across *MelonCore25* and QTL mapping in TAD×DUL RILs. **a)** Experimental design and examples of disease response of susceptible and resistant lines from 7 to 14 days post inoculation. **b)** Projection of the 25 lines response to FORC on the genetic PCA. Dashed line mark the cross (HDA034) used for the QTL mapping. **c)** Frequency distribution of FORC Disease Symptoms Index (DSI) across 153 TAD×DUL RILs. Response of parental lines is marked. **d)** Manhattan plot for linkage mapping of response FORC in the TAD×DUL RILs. **e)** Allelic effect of the chromosome 7 QTL. White line is the allelic mean. Gray boxes represent the standard deviation.

##### Variation in response to charcoal rot (Macrophomina phaseolina) across MelonCore25

We recently completed a multi-environment screen of the *MelonCore25* for resistance to the soil-borne fungal pathogen *Macrophomina phaseolina* causing a disease often referred to as charcoal rot (Cohen *et al*., 2022). The frequency distribution of cross-environment average disease severity index (DSI) of the 25 lines is plotted in **Sup. Figure 4a.** The whole spectrum from resistant to highly susceptible is represented across *MelonCore25*. Projection of the DSI values on the genetic PCA (**Sup. Figure 4b**) highlight the correlation between taxonomic classification (ssp. *melo* and *agrestis*) and resistance level, such that most *agrestis* lines show tolerance or moderate susceptibility, while most ssp. *melo* lines from the different variety groups show moderate or high level of susceptibility. Potential crosses are available between the tolerant and susceptible lines as part of the *HDA25F_2_s* set, and as such are good substrate for genetic dissection of this trait.

#### Unified framework for trait dissection in the pan-genomic era

As many horticulturally important traits are polygenic, the availability and utilization of multi-parental resources is required for comprehensive genetic dissection of natural variation and detection of repertoire of favorable alleles for crop breeding. Melon genetic and genomic research was enhanced since the release of the first melon genome 10 years ago (Garcia-Mas *et al*., 2012), followed by extensive re-sequencing of hundreds of melon accessions (Zhao *et al*., 2019; S., Liu *et al*., 2020; Dafna *et al*., 2021). Five additional high-quality melon genome assemblies that can be used as references for this species were published over the last 4 years (Zhang *et al*., 2019; Yang *et al*., 2020; Yano *et al*., 2020; Ling *et al*., 2021; Pichot *et al*., 2022). However, the recent expansion of the pan-genome concept, based on the realization that multiple genomes capturing the diversity within species are required for complete representation of the genomic variation (Li *et al*., 2022; Della Coletta *et al*., 2021; Bayer *et al*., 2020; Ho *et al*., 2020), suggest that the SV layer is not fully exposed in melon. We, therefore, developed and describe here a multi-parental trait dissection framework, anchored to a novel pan-genome constructed based on high-quality assemblies of 25 genomes, representing melon diversity. Pan-genomes are described for several crop plants, highlighting the commonality and magnitude of structural variation within species (Li *et al*., 2022; Qin *et al*., 2021; Y., Liu *et al*., 2020; Jayakodi *et al*., 2020; Song *et al*., 2020; Gao *et al*., 2019). The unique aspect of the current project is the creation of unified common framework that integrate between genetic diversity, phenotypic variation and genomic information. In addition to the traits described here, the *MelonCore25* set was phenotyped to fruit morphology traits (size and shape), rind and flesh color and pigments content (Gur *et al*., 2017), rind netting (Oren *et al*., 2020), fruit firmness and ripening behavior (Oren *et al*., 2022) and over 80,000 metabolomic and elemental features extracted from rind and flesh (Moing *et al*., 2020). Alongside the 25 genome assemblies and segregating populations covering all possible parental combinations, the genetic basis of these traits (and others) can now be studied more effectively. An important additional layer that can supplement the current framework is a comprehensive expression profiling (transcriptomes) of the *MelonCore25* across different organs, tissues and developmental stages, which will provide important layer for candidate genes discovery and will improve genomes annotations.

## Conclusions

We describe here a new resource for genomic research in melon. Pan-genome constructed from 25 diverse accessions selected to represent the broad variation across *Cucumis melo* is useful for comparative genomic analyses to address the evolution of melon genome and impact of breeding history. The integral connection of the pan-genome with a set of crosses and segregating populations (*HDA25F_2_*, n=300) is expanding the usability of this resource. For example, correlation between structural variation and parental crossability or fertility of F1s can be studied. Detailed analysis of relations between large genomic rearrangements and recombination frequency can be directly addressed using the SVs database and F2 populations. We demonstrate the primary potential of this integrated framework—genetic and genomic dissection of traits. The genomic information can also be very useful for informed design of genome editing and traits introgression in breeding programs, taking into consideration the impact of SVs.

## Materials and methods

### Plant materials and field trials

#### MelonCore25

This research is centered on the core set of 25 diverse melon accessions (**Table 1, Figure 1**) that were selected based on the comprehensive genotypic and phenotypic characterization of our broader *GWAS180* panel (Gur *et al*., 2017). The core set was selected based on multiple genotypic and phenotypic criteria as previously described (Dafna *et al*., 2021; Gur *et al*., 2017). Briefly, initial tentative set (n=40) was constructed to represent all the different horticultural groups in the diverse collection (based on traditional classification, (Pitrat, 2008)). Phenotypic profiles were then used as the second primary factor; the preliminary core set was projected on the distribution of the different traits to ensure phenotypic spectrum is well captured in the core panel (as illustrated at Gur *et al*. 2017). Following required adjustments and narrowing of the set to n=30, based on the first two steps, final set was selected to meet the 25 accessions target, taking into account maximum polymorphism information content (PIC) value and uniform distribution on genetic diversity plots.

#### HDA25

The creation of diverse, 25-way half-diallel population, resulting in 300 F1 hybrids that represent all possible parental combinations, was previously described by Dafna *et al*. 2021.

#### HDA20

a subset of HDA25, composed of 20 representative lines and their 190 half-diallel F1s. this subset is describe in Dafna *et al*. 2021.

#### HDA25F_2_s

All 300 *HDA25* F1s were grown in the greenhouse at Newe-Yaar and subjected to self-pollinations to produce the 300 F2 populations.

#### PI414□×□DUL RILs population (*HDA019*, F7)

A bi-parental population of 99 RILS from a cross between the mottled rind PI414723 (ssp. *melo* var. *momordica*) and the smooth rind parent Dulce (ssp. *melo* var. *reticulatus*) was previously described (Harel-Beja *et al*., 2010).

#### TAD□×□DUL RILs population (*HDA034*, F7)

TAD×DUL RILs population is composed of 164 F7 recombinant inbred lines originating from a cross between ‘Tam Dew’ (TAD; C. *melo* var. *inodorous*) and ‘Dulce’ (DUL; C. *melo* var. *reticulatus*) as previously described (Oren *et al*., 2020).

#### TAD × QME (*HDA008*) F6 population

The cross between the Honeydew line, TAD (ssp. *melo* group *Inodorus*) and the *cucumis colossus* line QME was grown at greenhouse at Newe-Yaar and ~200 lines were advanced to F6 through single-seed-descent selfing scheme.

#### TAD × PI164323 (*HDA192*) F5 population

the cross between TAD and PI164323 was grown at greenhouse at Newe-Yaar and ~150 lines were advanced to F5 through single-seed-descent selfing scheme.

#### SAS × DOYA (*HDA*243) F5 population

The cross between the sweet line SAS (ssp. *agrestis*, group *Makuwa*) and the non-sweet line DOYA was grown at greenhouse at Newe-Yaar and advanced to F5 through single-seed-descent selfing scheme.

All the above populations were grown in the open field at Newe-Yaar or in the greenhouse as previously described (Gur *et al*., 2017; Oren *et al*., 2020).

### Trait evaluation

#### Sugars analysis

Five mature fruits per plot were sampled in multiple harvests and total of 10-15 fruits per accession were analyzed. Concentrations of total soluble solids (TSS, measured in º Bx) were measured on flesh samples from each of the five fruits separately, using a hand-held refractometer (Atago Co. Ltd., Tokyo, Japan). Approximately one gram of frozen mesocarp tissue was placed in 80% EtOH and sugars were extracted and analyzed by HPLC as described (Petreikov et al. 2006).

#### Characterization of rind color and mottling pattern

rind traits were scored visually on young fruits on the vines at ~10-20 days post anthesis, and after harvest of mature fruit.

#### Response to fusarium wilt

protocols for pathogen and plant inoculation are described in details at Burger *et al*., (2003). *Fusarium oxysporum* f.sp*. melonis* used for the inoculation were maintained on yeast extract medium at 27 ° C in the dark. Conidial suspensions for seedling inoculation were prepared by macerating 1-week-old cultures with 100 mL water (Cohen *et al*., 1996). Seeds of *MelonCore25* were sown in sandy soil. Two days after emergence, the seedlings were removed from the soil and washed under running tap water. The roots were pruned to approximately half-length, and seedlings were inoculated by dipping the roots in a conidial suspension (10^6^ conidia mL^-1^) for 2 minutes. Inoculated plants were transplanted into 250 mL pots containing new, disease-free, sandy soil. Each tested genotype consisted of 35 seedling (five pots X seven plants per pot). First wilting symptoms were evident 7 days after inoculation. The number of wilted plants was recorded twice a week for two weeks and disease incidence was calculated.

#### FORC inoculation and symptoms scoring

Pathogen and plant inoculation are described in details at (Cohen *et al*., 2015). Briefly, melon seeds were sown in autoclaved sandy soil. Four days post emergence, seedlings were removed from the soil and washed under running tap water. The roots were gently pruned and the seedlings were inoculated by dipping the roots in a suspension of 10^6^ conidia mL^-1^ for 5 min. The inoculated seedlings were then transplanted into 250 mL pots containing clean sand and maintained in a growth room at a photoperiod regime of 12 h light/darkness. Plants were observed for 18 days following inoculation with FORC. First disease symptoms on ‘Dulce’ plants, leaf chlorosis, appeared six days after inoculation. Subsequently, the susceptible plants exhibited necrosis of the leaf margins, browning of the base of the stem, and wilting of the leaves and apical portion of the stem. The plant crown then became necrotic and the plant died. Eighteen days after inoculation, each plant was scored as symptomless (resistant) or as dead, wilted, necrotic or chlorotic (susceptible).

#### *Macrophomina phaseolina* inoculation and symptoms scoring

Pathogen and plant inoculation are described in details at (Cohen *et al*., 2022). Briefly, *M. phaseolina* (isolate no. NY 198) was isolated from melon plants. This isolate was tested routinely and is highly pathogenic. Melon plants at the age of 5 true leaves were inoculated with *M. phaseolina-* infested wooden toothpicks (Cohen *et al*., 2016). The plants were inoculated approximately 1 cm above ground level, by stabbing them with an infected toothpick and leaving it in the stab wound. Disease severity was evaluated based on lesion development and size on a 0-to-5 scale, with 0 = symptomless plant, 1 = initial small (< 1 cm) lesion, 2=1–3 cm lesion, 3 = stem rotting of 2–4cm, 4 = stem rotting of >5 cm, 5 = plant dead, completely necrotic. Accessions having average scores of ≤2.0 were considered to be resistant to *M. phaseolina* whilst those having average scores of≥3.0 were considered to be susceptible.

### Statistical analyses

JMP ver. 14.0.0 statistical package (SAS Institute, Cary, NC, USA) was used for statistical analyses. Mean comparisons were performed using the Fit Y by X function. GWA analysis was performed in TASSEL v.5.2.43 using the mixed-linear model (MLM) function. Distance matrix and Relatedness matrix of pairwise kinship (k matrix) were calculated in TASSEL from the filtered SNP dataset using the Centered_ IBS method (Endelman and Jannink, 2013). Stringent Bonferroni method was used to control for multiple comparisons in GWA.

### DNA preparation and Nanopore sequencing

High molecular weight (HMW) DNA extraction and size selection was carried out either using a modified CTAB protocol as previously described in Oren *et al*., (2022) or using Macherey-Nagel NucleoBond HMW DNA kit (REF 740160, www.mn-net.com). Library preparation was carried out following Oxford Nanopore Technologies (ONT) guidelines as described in Oren *et al*., (2022), and sequenced on MinION FLO-MIN106D flow cells. Base calling was done using the GPU version of Guppy (V5.0.11, Oxford Nanopore Technologies) using the HAC model.

### Genome assembly and scaffolding

ONT reads were assembled using Flye assembler v2.9 (Kolmogorov *et al*., 2019) by a previously detailed method (Oren *et al*., 2022), with the following adjustments: Following initial contig assembly, only one round of ONT based polishing was carried out using the built-in Flye polishing function, followed by one round of Illumina reads polishing using Pilon v1.24 (Walker *et al*., 2014). Reference guided scaffolding of contigs into pseudomolecules was carried out using RagTag v2.1.0 (Alonge *et al*., 2019) with the Harukei-3 as a reference (Yano *et al*., 2020). We used the Harukei-3 melon genome to avoid possible inverted assembly regions on the chromosome 6 of DHL92 as previously proposed (Yang *et al*., 2020).

### *Characterization of SVs* across *MelonCore25* and 300 *HDA25* crosses

Structural variants were characterized by whole genome alignment of each of the *MelonCore25* genomes to the DHL92 v4.0 genome using nucmer4 (Marçais *et al*., 2018) using –maxmatch –l 100 –c 500 parameters. The resulting delta file was then analyzed for InDels with command-line scripts of Assemblytics (v1.2.1) (Nattestad and Schatz, 2016), setting unique sequence length for anchors at 10kb. For short InDels database, InDel size was limited to a range of 15-2000bp—fragment sizes that can be effectively amplified using standard PCR assay. The Assemblytics output tables of all *MelonCore25* lines were concatenated, filtered to InDel type variants and splited by line to create a matrix of 194K InDels with a separate presence/absence column per line for each InDel. A simple conditional script was used to list the presence/absence of each InDel across the hybrids of the 300 nonreciprocal combinations of the 25 *MelonCore25* lines. The final InDel table is composed of 115,802 InDels, their attributes and 325 presence/absence columns (25 lines and 300 hybrids, **Sup. Table 3**).

### Validation of selected InDels and Genotyping of MT-2 and Fom-2 by PCR

Twenty-two InDels in size range of 16-1209 bp, that show polymorphism between DUL and PI414 were selected from our SVs database (**Sup. Table 4**, generated using Assemblytics (v1.2.1) (Nattestad and Schatz, 2016) for validation. Primers were designed using Gene Runner version 6.5. PCR, with an annealing temperature of 56°C, was performed on genomic DNA of Dulce, PI414 and their F1, using 2xPCRBIO HS Taq Mix Red (PCRBIOSYSTEMS, UK). The products were separated on a 2.5% or 1.2% agarose gel for 1-2 hours (**Sup. Figure 2, Sup. Table 4**). Primers for *MT-2* and *FOM-2* markers are listed in **Sup. Table 8.**

## Acknowledgements

We wish to thank Fabian Baumkoler and the Newe-Yaar farm team for technical assistance in setting the field trials and for plant maintenance. We appreciate and acknowledge Harry Paris for his critical language editing of the manuscript. Funding for this research was provided by the United States-Israel Binational Agricultural Research and Development Fund (BARD) grants no. IS-4911-16 and IS-5284-20.

## Author Contributions

### Conflict of interest

#### Data availability statement

The data supporting the findings of this study are available within the paper and within its supplementary materials published online. Raw sequences and FASTA files of genome assemblies can be found at NCBI [project information will be shared upon manuscript acceptance]

## Supporting information

**Supplementary Figure 1**: Workflow for using the pan-genome and multi-parental mapping framework.

**Supplementary Figure 2:** PCR validation of 22 InDels. InDel # correspond to **Sup. Table 4.**

**Supplementary Figure 3**: Mode of inheritance of TSS across the *HDA20* set: 190 hybrids and their parental lines.

**Supplementary Figure 4:** Variation in Disease Severity Index (DSI) to *Macrophomina phaseolina* across *MelonCore25*. **a**) Frequency distribution of DSI across *MelonCore25*. **b**) Projection of DSI on the genetic PCA. Diverse crosses are highlighted in dashed lines and hybrid code.

**Supplementary Table 1:** Nanopore long-read sequencing statistics.

**Supplementary Table 2:** Large SVs across *MelonCore25*.

**Supplementary Table 3:** Short InDels across *MelonCore25* and 300 *HDA25* crosses.

**Supplementary Table 4**: List of 22 InDels validated by PCR.

**Supplementary Table 5**: SNPs and short exonic polymorphisms within genes in the mottled rind interval (*MT-2*) on chromosome 2, across *MelonCore25*.

**Supplementary Table 6**: SVs within the mottled rind interval (*MT-2*) on chromosome 2, across *MelonCore25*.

**Supplementary Table 7:** Exonic polymorphisms in MELO3C021831 FOM-2 gene, across *MelonCore25*.

**Supplementary Table 8:** Primers for *MT-2* and *FOM-2* markers.

