## Supplementary Figures 1-5 for "Melon pan-genome and multi-parental framework for high-resolution trait dissection"

Sup. Figure 1: Workflow for using the pan-genome and multi-parental mapping framework

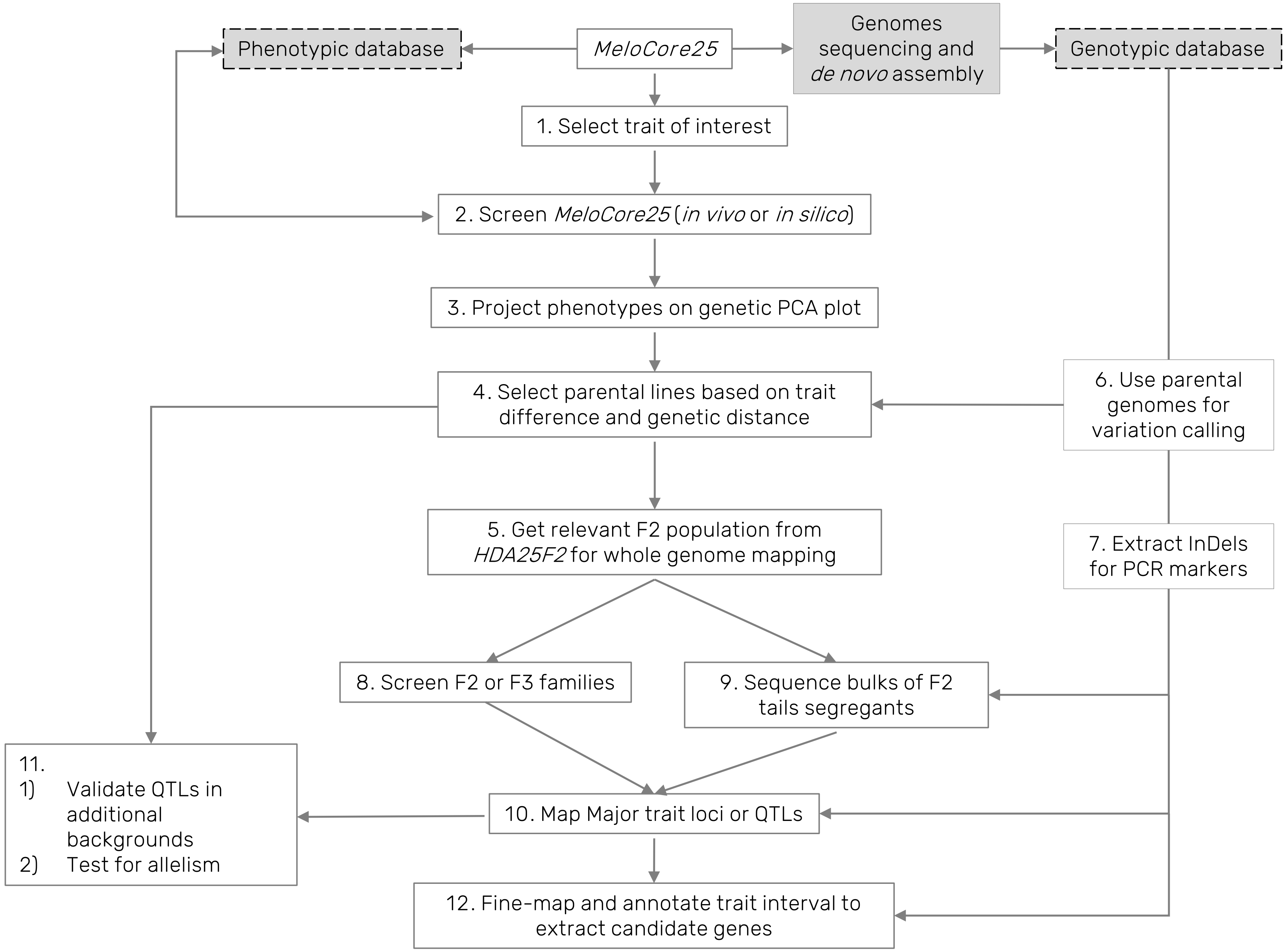

Sup. Figure 2: PCR validation of 22 InDels

A
2.5%
agarose

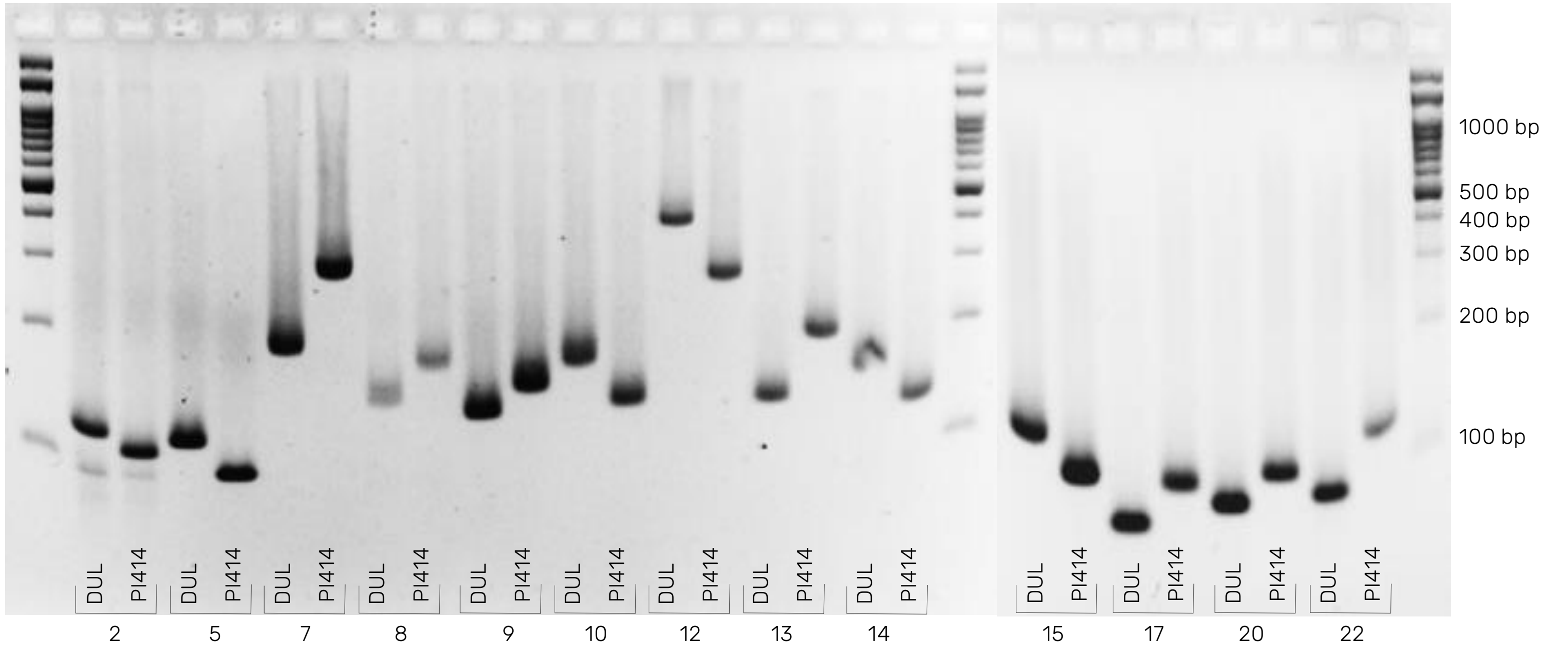


B
1.2%
agarose

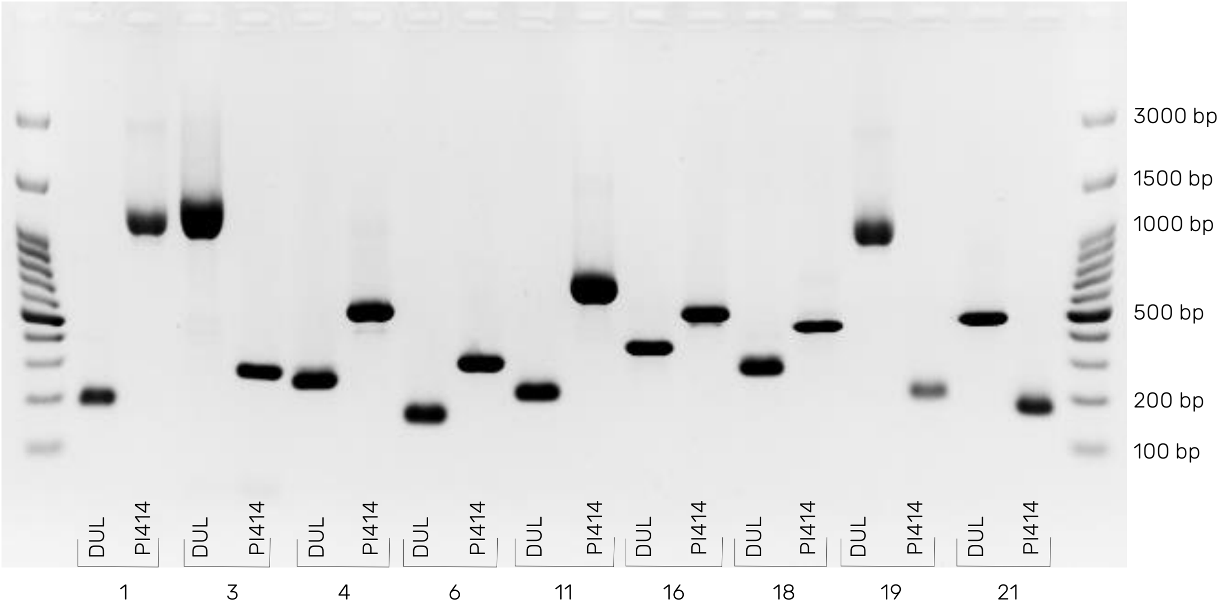

Sup. Figure 3: Inheritance of TSS across *HDA20* set: 190 hybrids and their parental lines

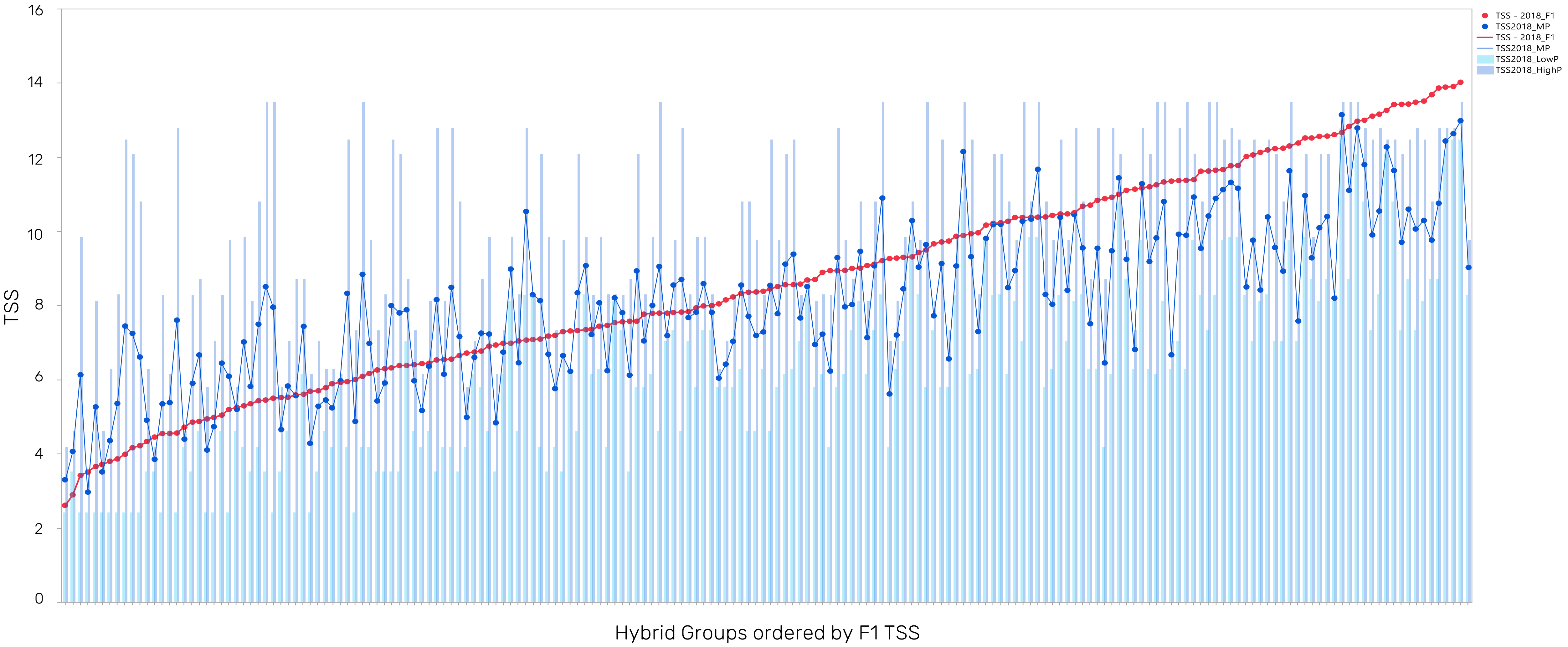

Sup. Figure 4: *Macrophomina* Disease response Index (DSI) across *MelonCore25*

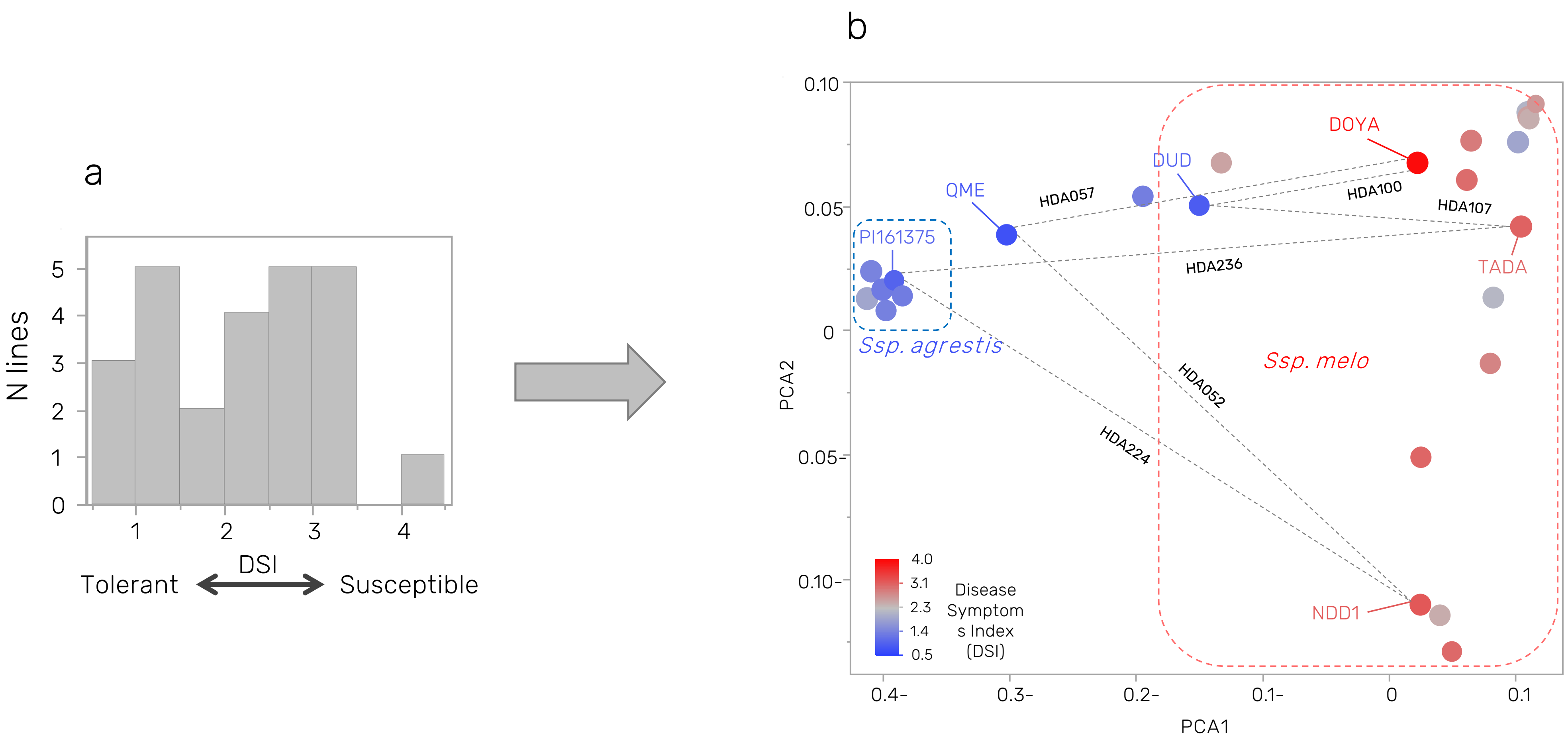

Sup. Figure 5: Comparisons between ONT and Ilumina reads alignments across 8 lines carrying the FOM-2 insertion.

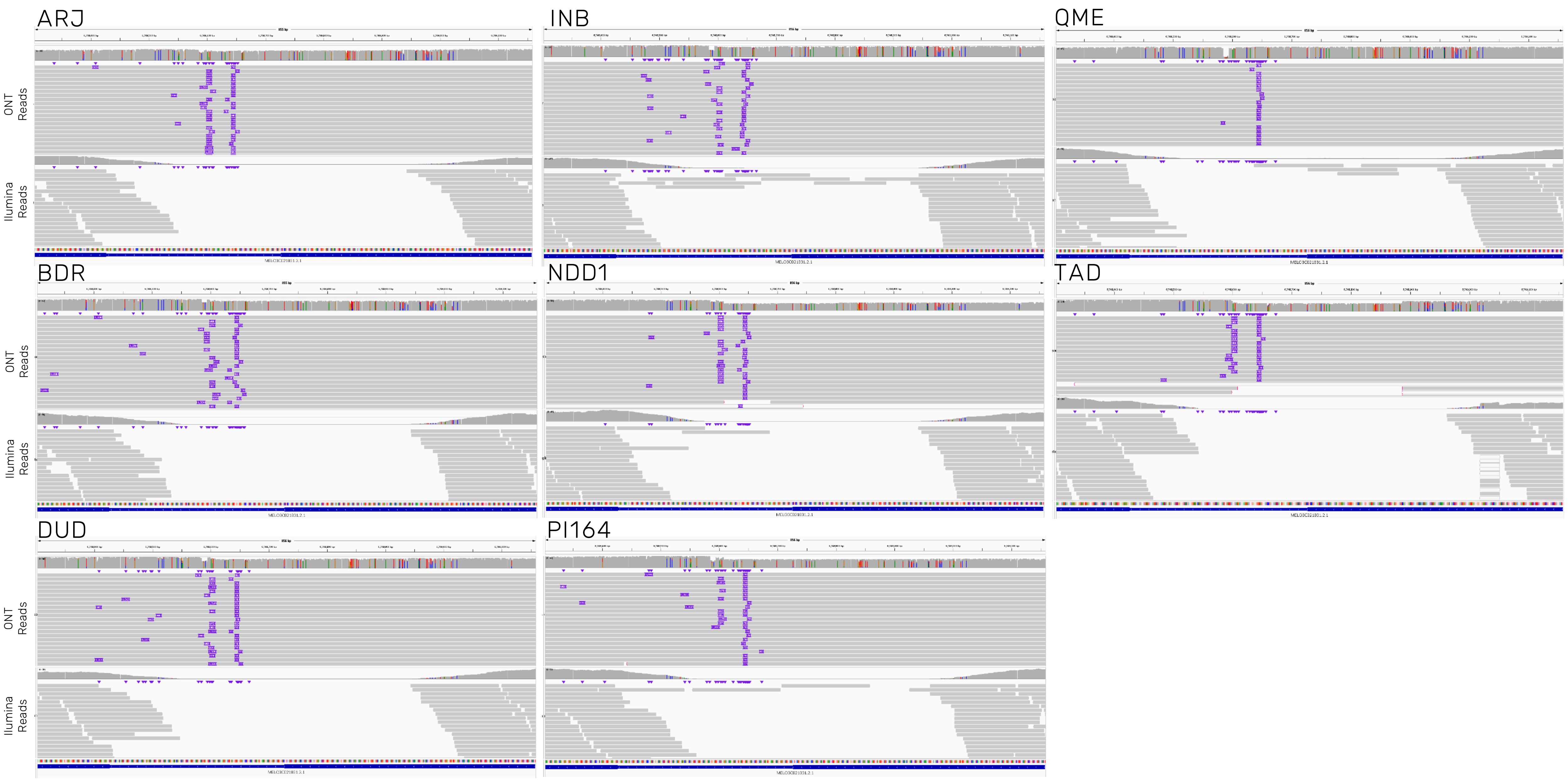
